# STEAP4^+^ neutrophils: a promising circulating biomarker for lung metastasis

**DOI:** 10.64898/2025.12.27.696660

**Authors:** Chen Ai, Huijuan Wu, Huihui Yang, Jing Wang, Chang Liu, Melissa S. F. Ng, Ziheng Zhou, Fenghui Zhuang, Qinhang Gao, Lingqian Wang, Changming Shi, Linnan Zhu, Zhaode Bu, Chang Chen, Lai Guan Ng, Zemin Zhang

## Abstract

As the leading contributor to cancer-related mortality, metastasis often evades detection until advanced stages, underscoring the critical need for early predictive biomarkers. Neutrophils are naturally circulating sentinels that have been implicated in pioneering pre-metastatic niche formation in distant organs. Nevertheless, their heterogeneity within evolving metastatic microenvironments remains poorly characterized. Here, through dynamically delineating neutrophil subsets via single-cell transcriptomics, we identified a conserved feature of lung metastasis in breast cancer and other cancer models. Among the core genes, a cell surface marker, STEAP4, was selected and validated to indicate lung metastases at both RNA and protein levels. In both humans and mice, these STEAP4^+^ neutrophils were detectable not only in lung metastatases but also in the circulation. Clinically, the abundance of circulating STEAP4^+^ neutrophils robustly discriminates patients with lung metastasis from those with localized primary tumor. Our findings uncover STEAP4^+^ neutrophils as a promising biomarker and lay the groundwork for a non-invasive, blood-based diagnostic strategy for metastatic disease.

## Introduction

Metastasis is the leading cause of cancer-related mortality, often diagnosed only at late stages, suggesting the critical need for early predictive biomarkers (Dillekas et al., 2019; Gerstberger et al., 2023; Siegel et al., 2023). The lungs are the most common sites of metastasis in many solid cancers (Altorki et al., 2019), and undergo early changes that create a favorable milieu for cancer cell colonization and metastasis initiation (Kaplan et al., 2005; Patras et al., 2023). This is evidenced by cellular and molecular characteristics of the lung metastatic tumour environment (TME) differing from those of primary-based tumours (Bohnenberger et al., 2018). Computed tomography (CT) can miss the diagnosis of lung micro-metastases at their early stages (Chen et al., 2019), and CTC-based detection exhibits low sensitivity due to their rarity in circulation (Elkholi et al., 2024). Hence, the ability to detect alterations in the lung (pre-)metastatic niche via biomarkers could facilitate early diagnosis of lung metastasis and improve treatment selection.

For a long time, neutrophils in both primary and metastatic tumors were viewed as a short-lived, homogenous population(Hedrick and Malanchi, 2022; Veglia et al., 2021b). Recently, however, increasing studies have demonstrated that they can differentiate into specific phenotypes to adapt to local tissue microenvironments, including the primary tumor microenvironment (TME) (Ballesteros et al., 2020; Ng et al., 2025; Ng et al., 2024; Palomino-Segura et al., 2023). However, whether neutrophils exhibit the same phenotypes in the metastatic niche as they do in the primary tumor remains poorly characterized. A key challenge in answering this question is that neutrophils in the metastatic niche may be influenced by organ-intrinsic property, signals directly from the primary tumor, or both.

To address this, we profiled neutrophil heterogeneity across steady-state, primary tumor and lung metastatic conditions, and determined a metastasis-specific gene program. We further pinpointed and validated STEAP4 as a cell surface marker for neutrophils bearing the metastatic gene program that was conserved in lung metastases across various cancer types. Remarkably, STEAP4^+^ neutrophils were detectable in the blood of metastatic patients. Their abundance could distinguish metastatic patients not only from healthy donors, but also from those with localized primary tumors. Our findings highlight that neutrophils adapt differently to primary and metastatic TMEs, resulting in different functional profiles that could be leveraged to develop a promising neutrophil-based biomarker for early detection of lung metastasis.

## Results

### Neutrophil accumulation coupled with deficient T cell activation discriminates lung metastases from primary lung tumors

Neutrophils, the most abundant immune cells in human blood, can infiltrate lung tumors, including both primary and metastatic lesions (Coffelt et al., 2015; Engblom et al., 2017; Salcher et al., 2022; Wculek and Malanchi, 2015; Zilionis et al., 2019). However, whether neutrophils exhibit functional heterogeneity in primary versus metastatic settings was understudied. To investigate this, we first compared neutrophil abundance in blood and lung tissues across healthy donors, patients with primary lung tumors (LUAD), and those with lung metastases (Table S1). Analysis of human peripheral blood revealed a significant increase in circulating neutrophils in breast cancer (BRCA) patients with lung metastasis, compared to the patients with primary lung tumors alone (Fig. 1A). In lung tissues, immunofluorescence staining further demonstated that heightened neutrophil infiltration in metastatic lesions relative to primary lung tumors or healthy lung tissue (Fig. 1B-C and Fig. S1A-B). These findings suggest that the elevated circulating neutrophils are strongly correlated with enhanced neutrophil recruitment to metastatic lung tumors, while showing minimal neutrophil infiltration in primary lung tumors or normal lung parenchyma.

**Figure 1.**
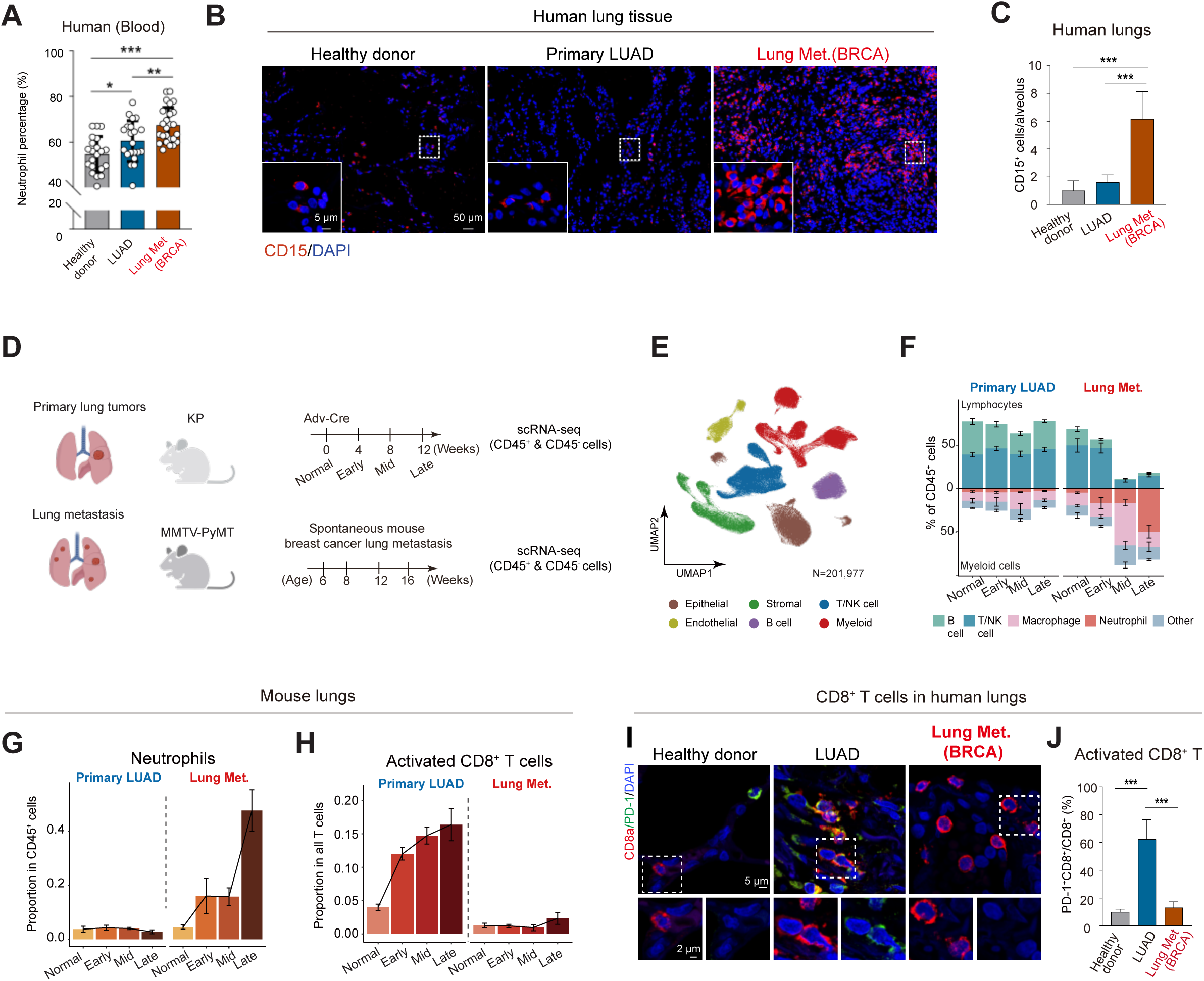
Neutrophil accumulation coupled with deficient T cell activation discriminates lung metastases from primary lung tumors. **A,** The proportion of neutrophils in whole blood from healthy donors, LUAD patients, and BRCA patients (mean ± SEM, n ≥ 20). \**p*<0.05, \*\**p*<0.01, \*\*\**p*<0.001, Student’s t test. LUAD: lung adenocarcinoma, BRCA: breast cancer patient. **B,** Typical images of immunostaining for CD15 in the lungs from the healthy donors, lung adenocarcinoma and breast cancer patients. Scale bars, 50 μm and 5 μm. **C,** The neutrophils labeled by CD15 in the alveolus were quantified based on immunostaining in **(B)**. Data: mean ± SEM, \*\*\**p*<0.001, Student’s t test. **D,** Experimental design for single-cell RNA-seq workflow. CD45^+^ and CD45^-^cells were sorted from mouse lungs of MMTV-PyMT and KP model at four time points for each condition. **E,** Uniform manifold approximation and projection (UMAP) plot of 201,977 cells from lung tissues of mice shown in MMTV-PyMT (n=108,387 from 15 mice) and KP models (n=93,590 from 13 mice). NK, natural killer cells. **F,** Mean of immune cell populations in KP and MMTV-PyMT mouse lungs as percentage of CD45^+^ cells. **G,** Bar plot showing mean neutrophil proportion in CD45^+^ cells across samples of each time point of KP and MMTV-PyMT models (mean ± SEM). **H,** Bar plot showing mean proportions of T cell subsets, activated CD8^+^ T cells (marked by *Pdcd1*) in KP and MMTV-PyMT mouse lungs. Mouse number of each time point in KP model: 4, 3, 3, 3; in MMTV-PyMT model: 3, 6, 3, 3 (Table S2). **I,** Typical images of immunostaining using antibodies against CD8a and PD-1 of lungs from LUAD, and BRCA patients. Scale bars, 5 μm and 2 μm. **J,** The percentages of CD8^+^PD-1^+^ T cells among CD8^+^ T cells in human donors are shown in (I). (mean ± SEM, n = 3). Student’s t test. \*\*\**p*<0.001.

The heterogeneity in neutrophil infiltration across lung tumors may reflect the divergent microenvironmental cues in primary versus metastatic lung TME. To dissect how the TME evolves differently in primary versus metastatic settings, we employed two murine models (Fig. 1D). For primary lung tumors, we utilized *Kras*^G12D^, *Trp53*-deficient (KP) mice to model lung adenocarcinoma that recapitulates the TME dynamics of human disease (DuPage et al., 2009) (Fig. 1D). For lung metastasis, we leveraged the MMTV-PyMT model, wherein mammary tumors spontaneously metastasize to the lung, enabling longitudinal analysis of TME remodeling during metastatic colonization (Guy et al., 1992; Lin et al., 2003) (Fig. 1D). To capture the fine-grained molecular landscape of TME evolution, we performed longitudinal single-cell RNA sequencing (scRNA-seq) across normal, early-, middle- and late-stage in both models(Fig. 1D; Materials and methods). Integration of the KP (n = 13) and MMTV-PyMT (n = 15) datasets generated a cross-disease lung tumor cell atlas, encompassing 201,977 high-quality cells identified spanning all stages(Fig. 1E; Fig. S1C; Table S2).

ScRNA-seq analysis of immune cell composition revealed a striking dichotomy between primary and metastatic lung tumors. While primary KP lung tumors maintained a lymphocyte-dominated TME, lung metastases in MMTV-PyMT mice exhibited progressive temporal evolution toward myeloid cell predominance (Fig. 1F). Notably, neutrophils accumulated in metastatic lungs as early as initial disease stages and became the dominant immune population in late-stage disease (Fig. 1G). In contrast, primary KP tumors showed marked proliferation of activated CD8^+^ T cells—defined by *Ifng*, *Gzmk*, and *Pdcd1* expression—while metastatic lesions maintained low CD8^+^ T cell frequencies (Fig. 1H). This distinct immune landscape was further validated by immunofluorescence staining: metastatic lesions displayed robust neutrophil infiltration (Fig. S1D-E), whereas primary tumors contained dense PD-1^+^ activated CD8^+^ T cell populations (Fig. S1F-G; (Ishida et al., 1992; Kansy et al., 2017). Together, these findings demonstrate that lung metastases develop a fundamentally distinct TME—characterized by progressive neutrophil dominance and static CD8^+^ T cell activity—compared to the lymphocyte-rich, CD8^+^ T cell-active environment of primary lung tumors, despite their shared anatomical location.

Conserved neutrophil accumulation was observed in lung metastases across both human and murine systems (Fig. 1B-C, S1D-E). To determine whether reduced T cell activation is associated with this myeloid-rich metastatic TME, we analyzed T cell states. In human patients, lung metastases consistently exhibited reduced expression of T activation signatures compared to primary lung tumors, even though lung metastases were originated from various cancer types (Fig. S1H-I and Table S3). Immunofluorescence staining further validated that metastatic lungs from BRCA patients harbored significantly fewer PD-1^+^ activated T cells compared to primary tumors (Fig. 1I-J). In mouse models, scRNA-seq profile of T cell states revealed that both CD4^+^ T cells and CD8^+^ T cells were enriched in naïve states in lung metastasis, compared to strong expression of activation-related genes in the early-stage of primary lung tumors (Fig. S1J-K). Furthermore, T cells in metastatic TME displayed reduced expression of interferon-γ (IFNγ) (Fig. S1L), while myeloid cells and stromal cells failed to upregulate the interferon response (Fig. S1L), reflecting impaired T cell activation in the lung metastatic niche (Gocher et al., 2022; Kaczanowska et al., 2021; Mellman et al., 2023). This aligns with recent studies demonstrating lung-infiltrating neutrophils robustly acquired potent immunosuppressive activity, including suppression of T cell proliferation and effector function in the lung metastatic niche (Gong et al., 2023; Wculek and Malanchi, 2015). Together, these data suggested that elevated neutrophil infiltration coupled with reduced T cell activation is a conserved feature of lung metastatic TME. The neutrophil accumulation, observed as early as metastatic initiation (Fig. 1G)(Wculek and Malanchi, 2015), may drive the establishment of an immunosuppressive microenvironment that is fundamentally distinct from the primary lung tumor, despite co-occurring in the same organ.

### Identification of a metastasis-specific gene signature in neutrophils within the metastatic niche

To identify neutrophil states unique to the metastatic lung microenvironment, our primary investigation focused on comparing neutrophils from primary lung tumors (KP model) with those from lung metastases (MMTV-PyMT model), thereby controlling for the organ niche, which was found to strongly influence neutrophil phenotypes(Ballesteros et al., 2020). A potential confounder in this approach, however, is the different tumor origins in the lung microenvironment. To address this, neutrophils from primary mammary tumors (Pri.Mammary) to served as the control for comparison with their corresponding lung metastases within the same MMTV-PyMT model (see Methods and Table S2). This intra-model analysis revealed markedly greater neutrophil heterogeneity in the metastatic site (Fig. 2A-C), confirming that the metastatic niche itself induces distinct neutrophil phenotypes, even within the same tumor genetic lineage. Furthermore, we observed that neutrophils in the MMTV-PyMT lung metastatic niche shared common subsets with those from normal lungs and KP primary tumors (e.g., the N3 subset), which were notably absent in the primary mammary tumor (Fig. 2C-D). This may be due to the lung tissue microenvironment exerting a terminal shaping effect on neutrophil phenotypes once they are recruited to the lung (Ballesteros et al., 2020; Gong et al., 2023; Palomino-Segura et al., 2023). Subsequently, we then employed the neutrophils from normal lungs and primary lung tumors as controls to exclude the lung-specific signature and isolate the metastasis-specific phenotypes in our subsequent analyses.

**Figure 2.**
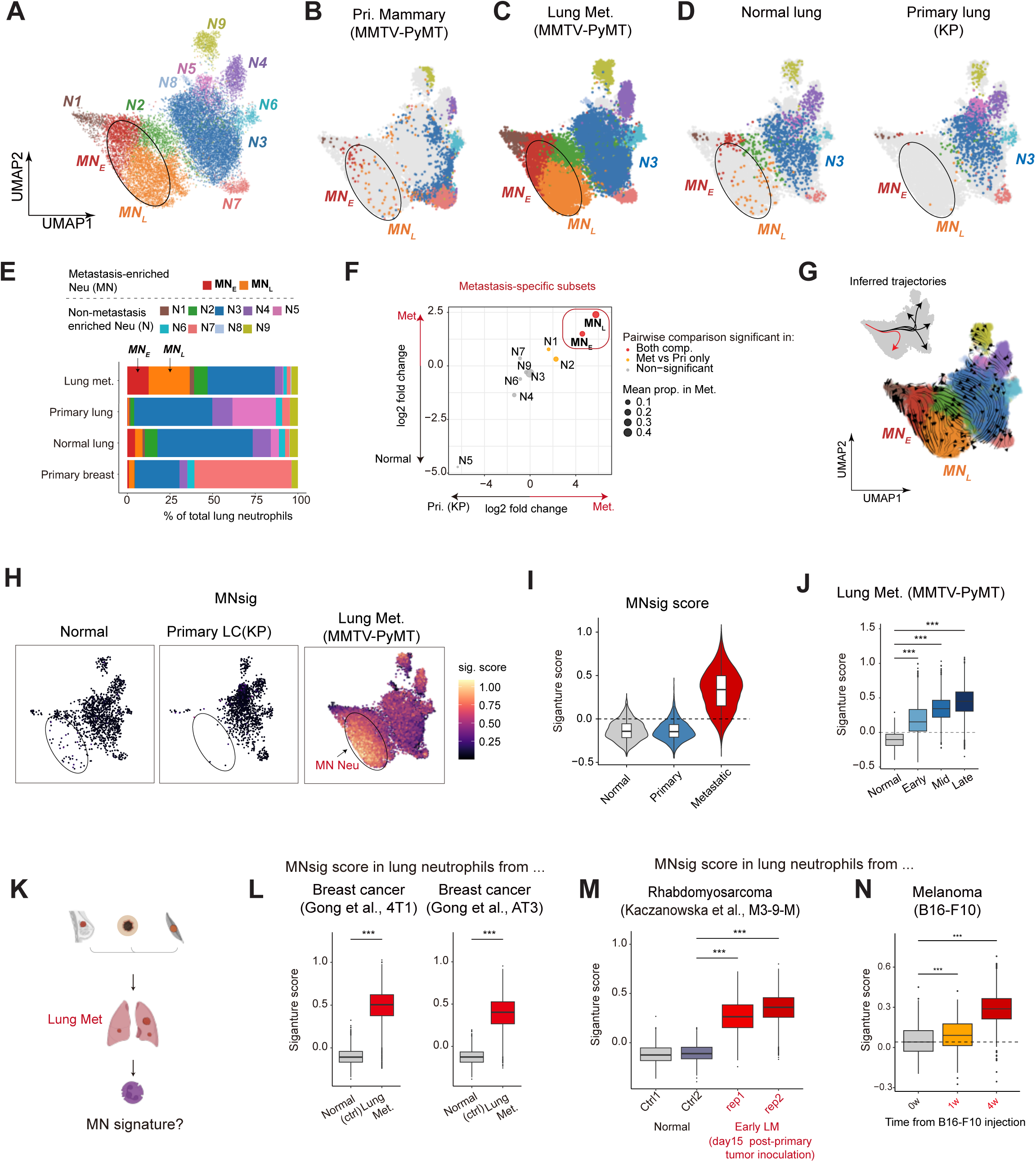
Neutrophils in the lung metastatic niche share a metastasis-specific gene signature across diverse tumor types. **A,** UMAP plots showing the neutrophil subsets integrated from B-D. **B-C,** UMAP plots showing neutrophil subsets in the integrated data of lungs from the primary sites and metastatic sites of the MMTV-PyMT model. Primary mammary tumor samples are from 12(mid) and 16 week(late) aged mice. **D,** UMAP plots showing neutrophil subsets in the integrated data of lungs from normal lungs and primary lung tumor from the KP model. **E,** Bar plot showing the proportion of each neutrophil subset within total neutrophils in each condition. **F,** Comparative analysis of neutrophil subset proportions in lung metastasis (Met.) versus normal lungs, and lung metastases in MMTV-PyMT versus primary lung tumors (Pri.) in KP model. Statistical Significance (*p*<0.05) was determined by the Wilcoxon rank-sum test. **G,** UMAP plots showing the direction of differentiation between neutrophil subsets inferred by RNA velocity and the Slingshot algorithm (left top). **H,** UMAP plots showing the expression level of MNsig in neutrophils from normal, primary tumor-bearing (from KP), and metastatic lungs (from MMTV-PyMT). **I,** Violin plot quantifying the distribution of MNsig scores in neutrophils across the three conditions, corresponding to the UMAP visualization in (H). **J,** Box plot showing the changes in score of MNsig signature genes (65 genes) across time within MN4 neutrophils from lung metastases in MMTV-PyMT. Student’s t-test. \*\*\**p*<0.001 **K,** Schematic of the analyses (shown in L-N) designed to test for the induction of the neutrophil MNsig in lung metastases from various tumor models. **L,** Comparison of MNsig levels in neutrophils from the normal lung and the metastatic niche, based on scRNA-seq data from (Gong et al., 2023). The analysis uses two orthotopic breast cancer mouse models, 4T1 and AT3. **M,** Comparison of MNsig levels in neutrophils from the normal lung and the metastatic niche, based on scRNA-seq data from (Kaczanowska et al., 2021) This analysis uses an orthotopic rhabdomyosarcoma mouse model at a pre-metastatic stage (day 15 post-primary tumor inoculation). **N,** Box plots comparing MNsig levels in neutrophils from normal lungs versus the metastatic niche, using scRNA-seq data from a B16-F10 experimental metastasis model created by tail vein injection.

Strikingly, we discovered two major populations exclusively enriched in the metastatic niche, Metastasis-enriched neutrophil Early (MN_E_) and Metastasis-enriched neutrophil Late (MN_L_) (Fig. 2A, 2C-F, Fig S2A). The MN_E_ and MN_L_ populations were virtually absent in all non-metastatic sites we examined: the primary MMTV-PyMT breast tumors, the primary KP lung tumors, and normal lung tissue (Fig 2A-F). Furthermore, the proportions of both populations progressively increased with metastatic advancement, while remaining absent throughout primary lung tumorigenesis (Fig. S2B). Taken together, these results indicate that the MN_E_ and MN_L_ populations are specifically induced by the metastatic context, rather than by the primary breast tumor or the lung organ niche.

To investigate the developmental relationship between MN_E_ and MN_L_, we performed a comprehensive analysis of their maturation and trajectory (Methods). A convergence of computational approaches confirmed a clear progression: RNA velocity analysis indicated a directional flow from MN_E_ to MN_L_(Fig. 2G), while trajectory inference with Slingshot modeled this as a distinct, metastasis-specific lineage (red, Fig. 2G). This differentiation progression was further supported by the ‘Neutrotime’ score, which positioned MN_E_ as an earlier subset, while MN_L_ represented a more mature, differentiated state (Fig. S2C). Collectively, these findings establish a clear MN_E_-to-MN_L_ differentiation trajectory that is unique to the metastatic niche.

The gene expression profiles of MN_E_ and MN_L_ were consistent with this differentiation axis and revealed a key distinction in their specificity. Consistent with their position as an intermediate state, MN_E_ expressed genes associated with early neutrophil maturation, such as *Prok2, Mmp8, S100a6,* and *Cd177* (Fig S2A). These markers, however, were not exclusive to the metastatic niche and were also detected in the early neutrophil populations within normal lungs and primary lung tumors(like N1 and N2) (Fig S2A), suggesting they include a general maturation gene program.

In contrast, the MN_L_ subset downregulated these early maturation genes(eg.*Cd177, Prok2*), and highly expressed genes such as *Steap4, Mif, Cstdc6*, and *Stfa2* etc., which were specifically enriched throughout the metastatic states but largely absent in non-metastatic contexts (Fig. S2A). We reasoned that as the stable, terminally differentiated endpoint of the MN_E_-to-MN_L_ trajectory, the gene program of the MN_L_ state would serve as the most robust representation of this metastasis-specific neutrophil identity. To validate this, we identify a gene signature from the top MN_L_ marker genes (Methods; Table S4), and observed the signature was highly specific to the lung metastatic niche, with negligible expression in normal tissues or primary tumors (Fig. 2H-I, Fig. S2D). Furthermore, differential expression analysis confirmed that this gene signature was robustly upregulated in the metastasis-specific populations (MN_E_ and MN_L_) relative to neutrophils from either primary lung tumors or normal lung tissue, highlighting its exclusive association with metastasis (Fig. S2E-F). Therefore, we defined these genes as the metastasis-specific neutrophil signature (MNsig) as a potential hallmark of metastasis-induced gene program (Table S4).

Notably, while most prominent in MN_L_ subset, MNsig was also expressed across other neutrophil subsets in the lung metastatic TME (Fig. S2A, S2G). This broad expression pattern suggests that MNsig represents a conserved gene program induced by the metastatic condition itself, rather than being a marker of one specific subset. Corroborating its link to disease progression, the overall expression of MNsig increased from early dissemination through advanced stages of metastasis (Fig. 2J). The signature’s high specificity and its dynamic expression during the metastatic cascade position it as a potential tissue-specific biomarker for lung metastasis.

The genes of the MNsig are enriched in metabolic and structural regulation. Their signature was characterized by genes involved in metabolic processes, particularly glycolysis and pyruvate metabolism, which are often upregulated to support the energetic demands of the metastatic cancer cells (Fig. S2H) (Welch and Hurst, 2019). Additionally, this signature was also enriched with cystatin genes (e.g., *Cstdc5*, *Stfa2*) linked to protease inhibition, which suggests MN neutrophils may modulate the extracellular matrix to promote metastasis.

### The MNsig is a conserved feature of neutrophils in the lung metastatic niche

To determine if the MNsig was a general feature of lung metastasis beyond our initial model, we investigated its expression in neutrophils from lung metastases derived from diverse primary tumors (Fig. 2K).

Breast cancer is a major cancer type that resulted in lung metastasis (Siegel et al., 2023). We first investigated that whether neutrophils from lung metastasis of other breast cancer models exhibit this signature. We analyzed public scRNA-seq data from two additional orthotopic breast cancer models, 4T1 and AT3 (Gong et al., 2023). In both models, neutrophils isolated from lung metastases showed elevated expression of the MNsig compared to those from normal lungs (Fig. 2L). Further corroborating this, analysis of bulk RNA-seq performed on neutrophils isolated from the lung metastatic niche of *K14*-Cre;*Cdh1*^fl/fl^;*Trp53^f^*^l/fl^ (KEP) mice also revealed a robust upregulation of the MNsig genes in a genetically engineered mouse model of spontaneous breast cancer (Fig S2I)(Garner et al., 2025). These data collectively establish that the MNsig is a conserved neutrophil signature within the lung metastatic niche across diverse breast cancer subtypes.

Next, we examined whether other cancer types induced metastasis-specific neutrophils in the lung metastatic niche. First, we analyzed the public scRNA-seq data from an orthotopic model of rhabdomyosarcoma (M3-9-M) that spontaneously metastasizes to the lungs (Kaczanowska et al., 2021). Notably, neutrophils within the lung already exhibited MNsig upregulation at a pre-metastatic stage (day 15 post-tumor inoculation) (Kaczanowska et al., 2021), suggesting this program is activated early during metastatic niche formation (Fig. 2M). This finding is consistent with our observations in the early-stage lung metastasis of MMTV-PyMT model (Fig. 2J).

Similarly, we utilized the B16-F10 experimental metastasis model, where intravenously injected melanoma cells rapidly colonize the lungs (Methods). Consistent with the previous reports, this model also led to an expansion of neutrophils in the lungs (Fig. S2J) (Hyun et al., 2020). Our scRNA-seq analysis of neutrophils harvested at 0-, 1-, and 4-weeks post-injection revealed a robust upregulation of the MNsig program as early as one week, which progressively increased through the fourth week (Fig. 2N). This temporal dynamic is highly consistent with the trend observed in the spontaneous MMTV-PyMT model, where the MNsig was also upregulated during the early stages of lung metastasis (Fig. 2J). Because this experimental model isolates the process of tumor cell colonization in the absence of a primary tumor, this observation suggests that the induction of the MNsig is not dependent on systemic signals from an established primary tumor, but is instead possibly induced locally in neutrophils within the lung microenvironment in response to tumor colonization. Together, the data from sarcoma and melanoma models indicate that the MNsig is not specific to a particular cancer type. Instead, it represents a conserved transcriptional program activated early in the metastatic stage by the lung microenvironment itself during the process of metastatic colonization. This suggests the MNsig may serve as a potential biomarker for lung metastasis originating from diverse cancer types.

### STEAP4 is a highly metastasis-specific membrane protein in both mouse and human metastatic niches

To advance the clinical translation of our findings on metastasis-related biomarkers, we aimed to identify a conserved cell surface protein from the MNsig gene program to enable practical clinical detection. Surface markers are prioritized over intracellular targets as they permit rapid, cost-effective analysis via flow cytometry—a widely available clinical tool. Crucially, an ideal marker must not only distinguish metastatic patients from healthy individuals but, more importantly, differentiate them from those with localized primary lung tumors. Current image-based diagnosis could fail to clearly discriminate between primary lung tumors and metastatic lesions, which disturbs clinical decision-making (Higuchi et al., 2021; Ichinose et al., 2016). Therefore, we compared the expression percentage of each gene in the MNsig in lung metastases of the MMTV-PyMT model, against those in normal lung tissues or primary lung tumors of the KP model (Fig. 3A). This dual-comparison strategy highlighted *Steap4* as the top candidate cell surface protein, exhibiting exceptional specificity for lung metastasis (Fig. 3B). *Steap4*, a metalloreductase involved in iron and copper homeostasis, may play an important role in the cellular response to inflammatory stress (Liang et al., 2017; Scarl et al., 2017; Xue et al., 2017).We next examined the expression of *Steap4* in our discovery dataset – the lung neutrophils of the KP and MMTV-PyMT models, and found that *Steap4* was highly specific to the metastatic niche, being broadly expressed across neutrophil subsets within the metastasis-bearing mice but absent in neutrophils originating from the normal lung or primary lung tumors (Fig. 3C).

**Figure 3.**
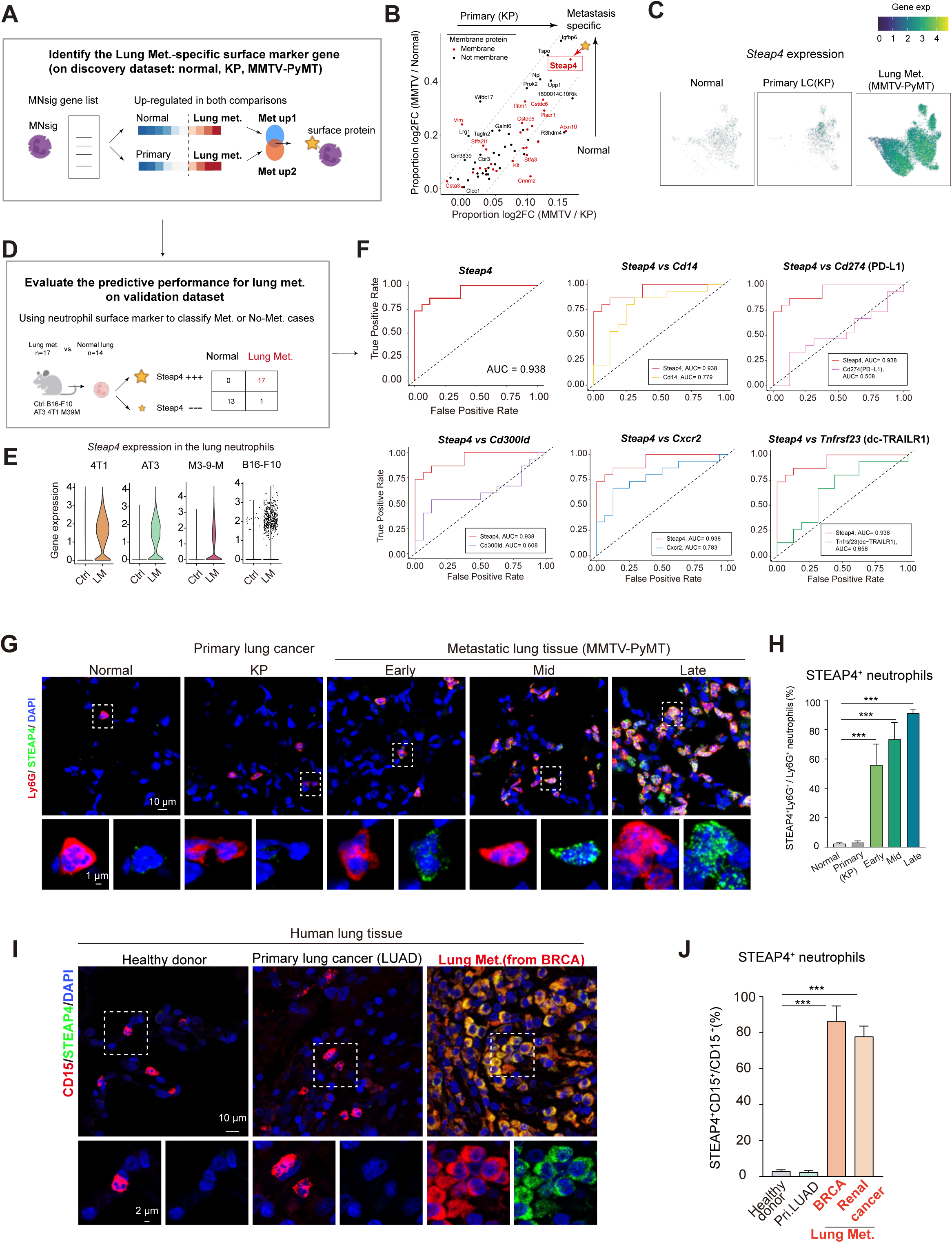
STEAP4, a membrane protein identified as a promising predictive marker for lung metastases. **A,** Strategy of identification of metastasis-specific surface protein marker genes in the discovery data from the MMTV-PyMT and KP models. **B,** Scatter plot depicting the metastasis specificity of genes in MN4 signature compared with normal lung tissue and primary lung tumor in KP. The proportion-weighted log2 fold change (log2FC) is calculated by combining the value of the mean expression proportion in neutrophils from MMTV-PyMT samples with the log2FC of the mean expression proportion across each group of samples. Membrane proteins are highlighted in red. **C,** UMAP plots showing the *Steap4* expression level in neutrophils from normal lungs, primary lung tumor and lung metastatic niche. **D,** Schematic depicting the approach for evaluating *Steap4* as a predictor for lung metastasis. The analysis incorporates validation datasets from B16-F10, AT3, 4T1, and M3-9-M models, compared against normal controls. **E,** Violin plot showing *Steap4* expression level across different lung metastasis models in the validation datasets in (D). **F,** The ROC curves measuring the performance of the expression percentage of each membrane protein on neutrophils in predicting lung metastasis using validation datasets in (D). **G,** Immunofluorescence staining using antibodies against Ly6G and STEAP4 of the lungs from normal, KP, and MMTV-PyMT (early, mid, late stage) models. Scale bars, 10 μm and 1 μm. **H,** Percentage of STEAP4^+^ neutrophils in total Ly6G^+^ neutrophils shown in (G). Student’s t-test. \*\*\**p*<0.001 (mean ± SEM, n = 3). **I,** Representative immunofluorescence staining of CD15 and STEAP4 in the lung tissues from healthy donors, LUAD patients and a BRCA patient. Scale bars, 10 μm and 2 μm. **J**, Percentage of STEAP4^+^ neutrophils among total CD15^+^ neutrophils in the sections from healthy donors and patients with lung metastases (mean ± SD, regions = 8, 8, 8, 4 for healthy donors, patients with lung cancer, breast cancer and renal cancer). \*\*\**p* < 0.001, Student’s t test.

To determine whether *Steap4* expression is a conserved feature of lung metastasis, we examined its expression in the cross-model validation dataset, including the 4T1, AT3, M3-9-M and B16-F10 datasets used above (Fig. 2L-N). Consistent with the pattern of MNsig, *Steap4* expression was uniformly up-regulated in neutrophils from the lung metastatic niche across all models(Fig. 3E). Furthermore, bulk RNA-seq data from neutrophils within the lung metastatic niche of KEP mice confirmed the upregulation of *Steap4* (Fig. S2I)(Garner et al., 2025). This remarkable consistency underscores its potential as a robust biomarker for lung metastasis.

Building on its potential as a biomarker, we next sought to quantify the predictive performance of *Steap4* for lung metastasis. We focused this analysis on the pre-clinical mouse models, given the inherent difficulty of studying the nascent pre-metastatic niche in patients (Fig. 3D). To do this, we pooled neutrophil data from the four validation models (Fig. 3D), creating a cohort of metastatic (n = 17) and normal (n = 14) lung samples. Although tumor-associated neutrophils expressed multiple immunosuppressive cell surface membrane proteins, their utility in distinguishing metastasis remains limited. These cell surface proteins include PD-L1 (Youn et al., 2008), CD14 (Veglia et al., 2021a), CD300LD (Wang et al., 2023), dcTRAIL-R1(*Tnfrsf23*)(Ng et al., 2024), CXCR2 (Acharyya et al., 2012; Chao et al., 2016), VISTA(*Vsir*) (Xu et al., 2019), and CD39(*Entpd1*) (Ryzhov et al., 2014). A direct comparison demonstrated that among these proteins, only *Steap4* consistently show high expression in lung metastases (Fig. S2K). Strikingly, ROC analysis revealed that *Steap4* exhibited unparalleled performance (AUC = 0.938), surpassing established markers like *Cd14* (AUC=0.78), and outperforming the known immunosuppressive genes like *Cd274* (PD-L1, AUC = 0.51), *Cd300ld* (AUC = 0.61). dcTRAIL-R1, VISTA, CD39(*Entpd1*), though enriched in tumor-associated neutrophils or macrophages (Ng et al., 2024; Ryzhov et al., 2014; Xu et al., 2019), showed no predictive value for lung metastasis (Fig. 3F and Fig.S2L). This finding was particularly notable as the cohort included samples from the pre-metastatic stage of the M3-9-M model (Kaczanowska et al., 2021) and B16-F10 experimental metastasis model, highlighting the sensitivity and robustness of *Steap4* as a biomarker.

To further validate STEAP4 on neutrophils at the protein level, we performed immunofluorescence staining on lung tissues from MMTV-PyMT mice. As predicted, STEAP4 was specifically expressed on mouse neutrophils within lung metastases, and staining was absent in normal lung tissues and primary lung tumors (Fig. 3G-H). Further, the proportion of STEAP4^+^ neutrophils emerged at the early-stage metastasis, and was significantly elevated as the lung metastasis progressed (Fig. 3G-H). These results demonstrate that the expression of STEAP4 is strongly associated with metastasis progression, positioning STEAP4^+^ neutrophils as a potential predictive marker for the occurrence of lung metastasis.

To determine whether STEAP4^+^ neutrophils were similarly specific to lung metastases in human patients, we performed immunofluorescence staining for STEAP4^+^ neutrophils in human lung tissue sections. In lungs from both healthy and LUAD patients, STEAP4^+^ neutrophils were rarely detected (Fig. 3I). In contrast, in the lung metastatic lesions from a patient with prior primary breast cancer, the proportion of STEAP4^+^ neutrophils within CD15^+^ neutrophils were significantly elevated, as observed in eight randomly selected regions (Fig. 3I-J). Similarly, lung metastasis in a patient with primary renal cancer also showed a striking increase in STEAP4^+^ neutrophils based on analysis of four randomly selected regions (Fig. 3J and Fig. S3A). Together, these findings demonstrate that STEAP4^+^ neutrophils uniquely emerge within lung metastatic niches in both mouse and human cancers, highlighting a conserved neutrophil response to lung metastases.

### STEAP4^+^ neutrophils as a promising blood-based biomarker for lung metastasis

Neutrophils are highly responsive and sensitive immune cells (Burn et al., 2021), making them uniquely suited to serve as early detectors of disease using immune cells. However, their potential as biomarkers for metastasis remains largely unexplored. Having identified STEAP4^+^ neutrophils within lung metastatic niches, we next investigated whether these neutrophils were confined to local metastatic sites, or could circulate in the blood—a feature that would make them a promising liquid-biopsy biomarker to assist in lung metastasis diagnosis.

To assess the presence of STEAP4^+^ neutrophils in the blood, we first analyzed *Steap4* expression in mouse models at the RNA level. Analysis of blood neutrophils via scRNA-seq demonstrated consistently elevated *Steap4* expression in the 4T1, AT3 and KEP mouse models of lung metastasis compared to neutrophils from either non-tumor bearing mice or a non-metastatic, primary tumor model of orthotopic pancreatic cancer (Fig. 4A and Fig. S3B) (Garner et al., 2025; Gong et al., 2023; Ng et al., 2024). These findings were corroborated by immunofluorescence staining of whole blood cells, which demonstrated that STEAP4^+^ neutrophils were uniquely detectable in blood neutrophils from mice bearing lung metastases. Importantly, these circulating STEAP4^+^ neutrophils emerged at the early-stage metastasis in the MMTV-PyMT model, with their frequency progressively increasing as metastasis progressed (Fig. 4B-C) – a pattern that mirrored the dynamics observed within the lung metastatic niche (Fig. 3G-H). This observation suggests STEAP4^+^ neutrophils show potential as an early-detection biomarker for lung metastasis.

**Figure 4.**
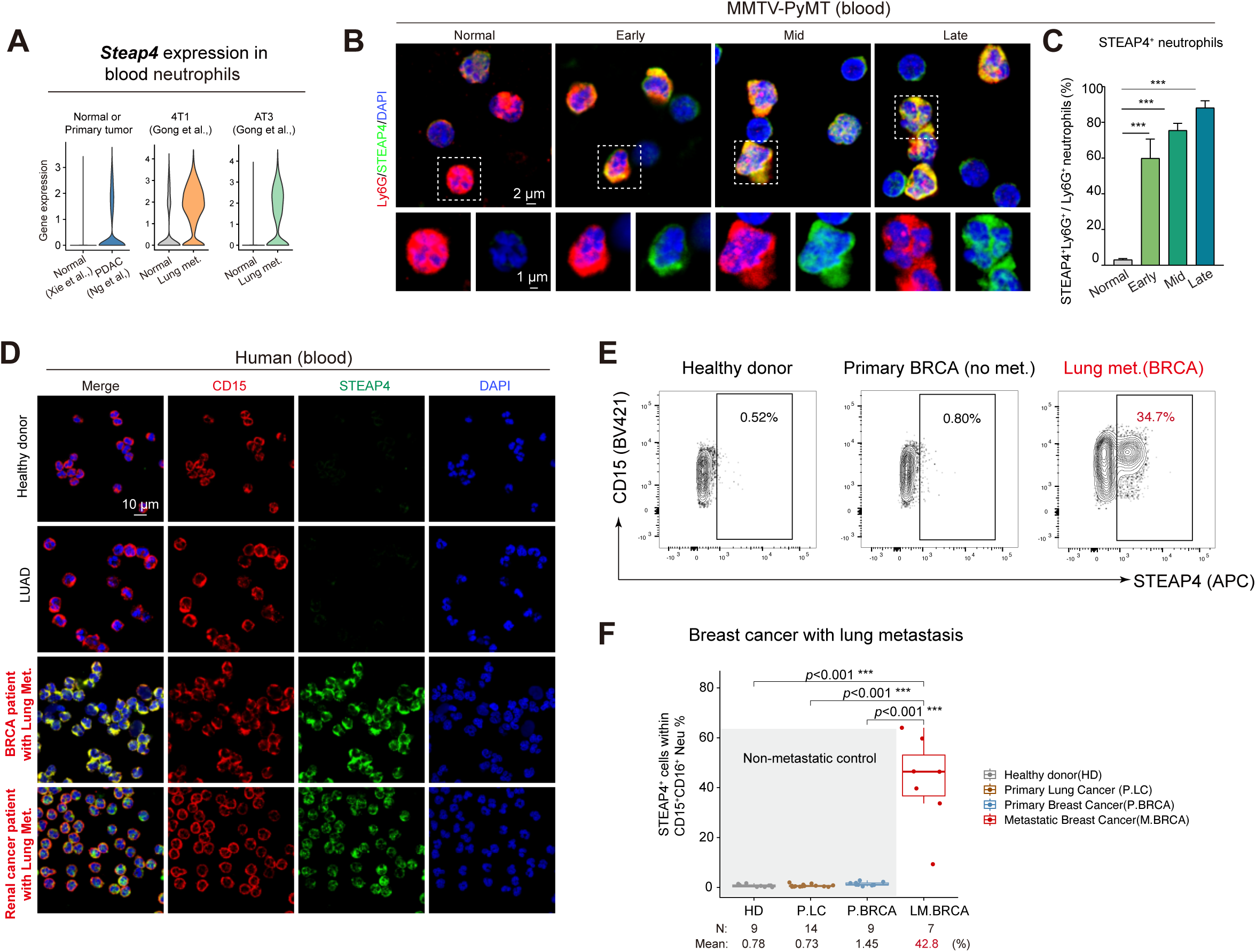
STEAP4^+^ neutrophils are uniquely elevated in peripheral blood of human cancer patients with lung metastases. **A,** Violin plot comparing *Steap4* expression in blood neutrophils from normal state, primary and metastatic states across different mouse models. **B,** Immunofluorescence staining using antibodies against Ly6G and STEAP4 of whole blood cells from normal, and MMTV-PyMT (early, mid, late). Scale bars, 2 μm and 1 μm. **C,** Percentage of STEAP4^+^ neutrophils in total Ly6G^+^ neutrophils shown in (**B**) Student’s t test. (mean ± SEM, n = 3). \*\*\**p*<0.001. **D,** Peripheral blood neutrophils from both healthy donors and lung metastasis patients were isolated and smeared. Representative immunostaining images of CD15, STEAP4 in the blood of healthy donor (n=10), LUAD patients (n=10), and patients with lung metastases (n=9). Scale bars, 10 μm. **E,** Representative flow cytometry scatter plots showing the percentage of STEAP4^+^ neutrophils in CD15^+^ blood neutrophils of healthy donors (n = 7), BRCA patients without metastasis (n = 4), and BRCA patients with lung metastases (n = 4). **F,** Box plot comparing the percentage of STEAP4^+^ neutrophils in human blood samples. Healthy donors (n = 9), patient with primary lung cancer (n=14), primary breast cancer (n=9), breast cancer with lung metastases (n=7). Student’s t-test. (Table S5)

Next, we examined whether STEAP4^+^ neutrophils could also be specifically detected in the blood of patients with lung metastases. We employed immunofluorescence staining on blood smears as a rapid method to assess the presence of these cells. The staining revealed that both healthy donors (n=10), and primary lung cancer patients (n=10) exhibit minimal STEAP4^+^ neutrophils in the blood. In contrast, lung metastases from multiple cancer types exhibited highly expressed STEAP4 protein on blood neutrophils (Fig. 4D and Table S1), despite their diverse tumor origins. These findings suggest that STEAP4^+^ neutrophils detected in the blood may mirror the metastatic remodeling occurring in lung parenchyma, underscoring that STEAP4^+^ neutrophils may serve as a promising blood-based biomarker capable of discriminating patients with lung metastases from both healthy and those with primary lung tumors.

We next employed flow cytometry on fresh blood samples to quantitatively investigate whether STEAP4^+^ neutrophils exhibit a distinct abundance in breast cancer patients with lung metastases, compared to non-metastatic breast cancer patients, primary lung cancer patients and healthy donors (Table S5). Neutrophils were selected and stained with anti-CD15 and anti-CD16 antibodies (Fig. S3C). In healthy donors (n=9), STEAP4^+^ neutrophils constituted a rare subset of neutrophils (mean 0.78%) (Fig. 4E-F and Fig. S3D). A similar low proportion was observed in patients with non-metastatic primary tumors, including lung cancer (mean 0.73%; n = 14), breast cancer (mean 1.45%, n=9). In contrast, the proportion of circulating STEAP4^+^ neutrophils was significantly elevated in the breast cancer patients with lung metastases, which accounted for a mean of 43% (n=7) of the neutrophil population—a nearly 30-fold increase over patients with primary breast cancer (Fig. 4E-F and Fig. S3D).

## Discussion

In summary, our integrated pre-clinical and clinical analysis identifies STEAP4⁺ neutrophils as a novel biomarker for lung metastasis. We validated this in fresh human blood samples, finding that circulating STEAP4⁺ neutrophils are significantly abundant in breast cancer patients with lung metastasis, but not in healthy donors, primary breast cancer patients, or patients with primary lung cancer. This discovery highlights the potential for a STEAP4-based blood assay, offering a non-invasive tool to detect lung metastasis and holding significant translational value for improving patient survival.

The translational potential of this finding is further supported by the intrinsic properties of neutrophils: (1) they are among the first responders to metastatic changes (Szczerba et al., 2019; Wculek and Malanchi, 2015); (2) they constitute a majority proportion of circulating immune cells, and (3) their high adaptability makes them highly sensitive to metastatic changes. Our data in MMTV-PyMT mice demonstrate that STEAP4^+^ neutrophils are detectable as early as 8-week age, suggesting their utility in early metastasis detection (Fig. 3G-H and Fig. 4B-C). Current liquid biopsy approaches for detecting tumor metastasis predominantly rely on biomarkers derived from primary tumors, such as circulating tumor cells (CTCs) (Elkholi et al., 2024). However, these methods face critical limitations: CTC-based detection exhibits low sensitivity due to their rarity in circulation and heavily depends on tumor-specific mutation profiles for identification (Elkholi et al., 2024). In contrast, neutrophils are more abundant in peripheral blood and are independent of tumor origin or tumor mutational profiles. Our findings highlight the potential of a STEAP4-based blood assay as a non-invasive tool for metastasis prediction and detection. This establishes the neutrophil-based signature as a novel avenue—distinct from CTC detection—that holds substantial translational value. Therefore, we propose that STEAP4^+^ neutrophils may serve as a sensitive, blood-based biomarker for the early monitoring of metastasis risk.

This discovery aligns with the established concept that neutrophils acquire context-specific gene programs to adapt to different pathological conditions (Ng et al., 2025). While both anti-tumoral and pro-tumoral neutrophil subsets have been described in primary tumors (Engblom et al., 2017; Ng et al., 2024; Wu et al., 2024), the gene programs specific to metastasis remain poorly understood. Our findings help bridge this gap by identifying the neutrophil gene signature specific to lung metastasis. We acknowledge that our study is currently limited by the lack of functional experiments regarding STEAP4^+^ neutrophils. Nevertheless, given the substantial clinical significance of this discovery, we believe it is critical to report these findings to provide a roadmap for future mechanistic and therapeutic studies.

## Materials and methods

### Human subjects

All procedures involving study participants were approved by the Human Research Ethics Committee of the Shanghai Pulmonary Hospital (K24-581). Clinical information of all subjects in this study was listed in Supplementary Table S1.

### Computed Tomography (CT)

All the donor and patients underwent thin-section CT. All CT examinations were performed with a 64-section scanner (Somatom Definition AS+, Biograph64), Philips (Brilliance 40, iCT256, Ingenuity Flex, MX 16-slice) with contrast material, and United Imaging (uCT 510, uCT 760, uCT S-160). Chest CT lesions in each patient were identified by pathologists.

### Histology for human lung tissues

Human Adult Lung tissue samples were obtained from LUAD and breast BRCA patients at the time of surgery. Additionally, normal lung tissue from adult donor lungs unsuitable for transplant and lung adenocarcinoma tissues from lobectomies were included. Fresh tissues were fixed with 4% paraformaldehyde (PFA) overnight at 4°C, and paraffin sections (4 µm) were prepared for immunofluorescent (IF) analysis and Hematoxylin and Eosin (H&E) staining.

For the H&E staining experiment followed as our previous study.(Wu et al., 2020) Briefly, slides were deparaffinized and rehydrated. The nuclei were stained with hemotoxylin (Abcam, ab150678) for 2 minutes and the cytoplasm were stained with eosin (Sigma, HT110280) for 3 minutes. Slices were then sealed with neutral resin after the dehydration and clearing.

For IF, slides were deparaffinized and rehydrated, then washed 3× for 5 minutes in PBS. A ring was drawn around the tissue using a hydrophobic pen (Vector, cat. no. H-4000). Slides were washed 310 min in PBS. Tissue was permeabilized with PBS-0.1%TritonX-100 (Sigma-Aldrich, cat. no. T9284-100 mL) for 10 min. Next, slides were washed 310 min with PBS. The slides were incubated with blocking buffer (3%BSA (biofroxx, 4240GR100) + 0.5% Tween-20 + 10% normal donkey serum (Servicebio, cat.no. G1217-5ML)) filtered through a 0.22-um filter, for 1 hour in a humidified chamber. Primary antibodies (rabbit anti-CD8a, Abcam, ab237709; mouse anti-PD-1, Servicebio, GB12338; mouse anti-CD15, Abcam, ab241552; mouse anti-CD66b GeneTex GTX19779; rabbit anti-CD16 Abclonal, A23541; rabbit anti-STEAP4, Proteintech,11944-1-AP) were diluted in antibody dilution buffer (Ai Fang Biological, AFIHC014) overnighter in a humidified chamber on a shaker at 4C. Slides were then washed 3 10 min with PBS-0.1%Tween-20 (Sangon Biotech, cat. no. A600560-0500), sections were stained with matched secondary antibodies (Alexa Fluor 647 Donkey anti-mouse, Jackson Immuno Research, 715-605-151; Alexa Fluor 568 Donkey anti-rabbit, Thermo Fisher Scientific, A10042; Alexa Fluor 647 Donkey anti-rabbit, Jackson Immuno Research, 711-605-152) at RT for 2 h. Staining with secondary antibodies alone was used as the control. The slides were washed 610 min with PBS-0.1%Tween-20, and a final wash was performed with PBS. Slides were mounted by adding 20 uL Antifade Mountain containing 4’, 6-Diamidino-2-Phenylindole, Dihydrochloride (DAPI) (Beyotime, cat. no. P0126-25) to each tissue. Then a coverslip was carefully placed on top of the slides. Fluorescence images were acquired using a confocal microscope (Olympus FV4000). All the images were further processed with OlyVia software.

### Comparison of T cell genes and signatures between primary and metastatic tumors in patient cohorts

Clinical data on The Cancer Genome Atlas (TCGA) was downloaded by UCSCXenaTools, and gene expression matrix for LUAD TCGA RNA-seq samples were retrieved from the supplementary data of Zheng and colleagues (Zheng et al., 2019). META-PRISM data was retrieved from supplementary data of Pradat and colleagues, in which only lung metastases originating from extrathoracic cancers in the lung tissue were included (Pradat et al., 2023). Both expression data were processed by the same pipeline using Kallisto+TxImport and then transform by log2(TPM+1) for further analysis. To correct the unwanted technical variates between samples after merging samples from two cohorts, we employed RUV-III-PRPS (Molania et al., 2023), a method suitable for integrate and normalize multiple sources of transcriptomic datasets, to remove the variation caused by library size, tumor purity and batch effects from TCGA and META-PRISM cohorts. Beriefly, RUV-III-PRPS method is a linear model through which the presence and impact of unwanted factors can be inferred via technical replicatesand negative control genes. These technical effects were further removed from the expression data, to obtained the corrected expression data for downstream comparison analysis. In this process, tumor purity were calculated by ESTIMATE (Yoshihara et al., 2013). To mimic technical replicates, pseudo-samples with the same biology setting but different technical variates(e.g. tumor purity) were created. To estimate the difference caused by batch effects of two cohorts, we included LUAD tumors in META-PRISM as control data. Negative control genes for RUV-III-PRPS were identified as genes that were not affected by the cancer types, but highly affected by cohort datasets (within LUAD patients), tumor purity (within each cancer type from each dataset), and library size. Gene expression data that were removed from technical effects was obtained by RUV-III linear model for further comparison.

Eight T cell signatures involved with T cell states and recruitment from Bagaev and collegues were used in the analysis (Supplementary Table S3) (Bagaev et al., 2021). GSVA methodwas applied to calculate the signature score for each sample, with mean score of each group of samples were showing in the heatmap Fig. S1 (Hanzelmann et al., 2013).

### Establishment of mouse lung tumor models

The establishment of mouse lung tumor models was based on established methods outlined in the previous work (DuPage et al., 2009). Specifically, 2-month-old KP mice (B6-KP, GemPharmatech, Strain NO. T015831) were chosen and performed anesthetic treatment. Then, an endotracheal cannula was inserted into the trachea to deliver adenoviral viruses expressing Cre or scramble shRNA into the mouse lungs (Obio Technology company, Shanghai, China). Each mouse received a dosage of 2.5 x 10^7 viral genome copies diluted in warm 50 ml sterile saline. After treatment, mice were kept on a warm pad until they recovered, with lung tissues collected at different time points (0, 4, 8, 12 weeks) post-viral delivery for single-cell sequencing(Table S2).

Spontaneous breast cancer lung metastasis mouse model, MMTV-PyMT mice (FVB-MMTV-PyMT, GemPharmatech Strain, NO. T004993) were established using female mice obtained from GemPharmatech, known to develop mammary gland carcinoma and progressive lung metastasis from 6 weeks of age onwards. Following housing in a controlled environment, lung tissues were collected at different time points (6-, 8-, 12-, 16-week age) for single-cell sequencing. In addition, breast tissues were harvested at 12 and 16 weeks from two mice whose corresponding lung tissues were included in the analysis, allowing for matched inter-organ comparison (see Table S2).

To establish melanoma lung metastasis models, B16-F10 cells(B16-F10, ATCC, reference CRL-6475) were cultured and prepared for mouse modeling(mouse: C57BL/6, Charles River Laboratories, reference C57BL/6NCrl). After two passages in a cell culture incubator, cells were injected into two-month-old female mice via the tail vein. Each mouse received a slow intravenous injection of 100,000 cells in a total volume of 100 µL. Subsequently, lung tissues were collected at different time points (0, 1 and 4 weeks) after the above injection for single-cell sequencing.

All animals were housed in specific pathogen-free (SPF) conditions at the Peking University animal facility, adhering to ethical regulations. All animals were experimentally naive, and all procedures were conducted in accordance with national guidelines for laboratory animal housing and care, as well as institutional regulations reviewed and approved by the Institutional Animal Care and Use Committee at Peking University.

### Histology for mouse lung tissues

Mice were anesthetized and right cardiac perfused with PBS. Lungs were inflated with 4% paraformaldehyde (PFA) and were continually fixed in 4% PFA at 4°C for 24 hours. Then the lungs were cryoprotected in 30% sucrose and embedded in optimal cutting temperature (OCT, Tissue Tek). H&E was performed as detailed above. For immunofluorescence, the lung tissues were cryosection at 15-μmthickness and air-dried overnight at RT. After removing the OCT, Sections were blocked and permeabilized with 3% bovine serum albumin (BSA) plus 0.1% Triton X-100 in PBS for 1 hour, and then incubated overnight at 4°C with primary antibodies (rabbit anti-CD8a, Abcam, ab217344; mouse anti-PD-1, Servicebio, GB12338; rat anti-Ly6G, Biolegend, 127602; rabbit anti-STEAP4, Proteintech, 11944-1-AP). After washing, sections were stained with matched secondary antibodies (Alexa Fluor 647 Donkey anti-mouse, Jackson Immuno Research, 715-605-151; Alexa Fluor 568 Donkey anti-rabbit, Thermo Fisher Scientific, reference A10042; Alexa Fluor 647 Donkey anti-rat, Jackson Immuno Research, reference 712-605153; Alexa Fluor 647 Donkey anti-rabbit, Jackson Immuno Research, 711-605-152 at RT for 2 h. Images were captured with a confocal fluorescence microscope (Olympus, FV4000).

### Single cell collection, library preparation, and sequencing for mouse samples

After deep anesthesia, mice at respective time points underwent subsequent procedures. Specifically, to eliminate blood cells from the lungs, mice were perfused with phosphate-buffered saline (PBS) through their left ventricles. Subsequently, a 1 mL enzyme solution comprising 5 U/mL neutral protease (Worthington-Biochem, LS02111) and 0.33 U/mL DNase I (Roche, 10104159001) was instilled into the lungs via the trachea. The lungs filled with the enzyme solution were further immersed in the same solution for 45 minutes at room temperature. Following enzymatic incubation, the digested lung tissues were gently minced into small pieces and incubated in Dulbecco’s modified Eagle medium 1640 (Invitrogen) containing 10% FBS for 10 minutes at room temperature. A single-cell suspension was obtained by passing the cells through 100 mm and then 40 mm cell strainers. The cell suspension was subsequently centrifuged at 800 rpm for 10 minutes to form a pellet. This pellet was resuspended in an RBC lysis buffer and incubated at room temperature for 10 minutes to eliminate red blood cells. After centrifugation at 800 rpm for 10 minutes, the supernatant was removed.

Next, cells were resuspended in FACS staining buffer to a final volume of 1 mL, and anti-CD45 antibody (PE/Cyanine7 anti-mouse CD45 Antibody, biolegend, 103114) was added at a ratio of 1:500. The cells were gently agitated every 10 minutes to prevent cell clumping and settling. After 30 minutes, unbound antibodies were removed using FACS staining buffer, and the centrifuged cells were resuspended in fluorescence-activated cell sorting (FACS) buffer prior to DAPI staining. CD45^+^ cells were sorted using the single-cell select mode in BD FACS Aria II and III instruments.

Subsequently, an equal number of CD45^+^ and CD45^-^ cells were mixed in a 1:1 ratio and processed following the 10X genomics protocol. Briefly, sorted cell subsets were loaded onto the 10X Chromium system and encapsulated using the Single Cell 50 Library & Gel Bead Kit (Chromium Single Cell 50 Library and Bead Kit, 10X Genomics, cat.no. 1000006; Chromium Single Cell 50 Library Construction Kit, 10X Genomics cat.no. 1000020). Single-cell gene expression profiles were generated according to the manufacturer’s instructions. The completed libraries were sequenced on HiSeq4000 Illumina platforms with a targeted median read depth of 50,000 reads per cell from total gene expression libraries.

### Neutrophil isolation from human whole-blood samples

Fresh whole-blood samples were collected from human donors. Peripheral blood was slowly layered onto Polymorphprep (Polymorphprep,No.1895) without mixing the blood into the lower Polymorphprep layer. Centrifugation was performed at 500 g for 35 min at 20°C with the lowest acceleration/deceleration settings.The neutrophil layer (polymorphonuclear leukocytes) was carefully aspirated using a pipette, avoiding contamination from the upper mononuclear cell layer. Cells were washed with 5 mL of 50% HBSS (Gibco,24020117) and resuspended by gentle pipetting. Subsequent centrifugation steps included: 350 g for 10 min at 20°C, followed by supernatant removal (avoiding cell loss by not tilting the tube). Cells were resuspended in 10 mL HBSS, centrifuged at 300 g for 5 min at 20°C, and treated with RBC lysis buffer (Invitrogen, 00-4333-57). After lysis, cells were centrifuged again (300 g, 5 min, 20°C), washed with 10 mL HBSS, and pelleted at 300 g for 5 min at 20°C. The final pellet contained purified neutrophils.

### Flow cytometry

Isolated neutrophils were resuspended in PBS containing 2% FBS and then incubated with FcR blocking reagent human (BD,564220) for 15 min. Cells were incubated with fluorochrome-conjugated antibody mix (BD Horizon™ BV421 Mouse Anti-Human CD15,BD 567008; Alexa Fluor® 647-conjugated Anti-Human STEAP4, R&D Systems FAB4626R; BD Pharmingen™ PE Mouse Anti-Human CD16, BD 561313) for 30 min at 4°C in the dark. Cell suspension was washed twice and stained with eBioscience™ 7-AAD Viability Staining Solution (eBioscience,00-6993-50) and analyzed on a BD FACS Aria SORP. Data processing was performed using FlowJo™ v10 software (FlowJo LLC).

### Immunostaining of whole-blood samples

Whole-blood samples from normal mice, MMTV-PyMT (early, mid, late) mice were collected in heparin-containing collection tubes. After two rounds of red blood cell lysis with 1x RBC lysis buffer (Invitrogen, 00-4333-57) for 5 minutes and wash with working buffer (5%FBS+PBS), the single-cell suspensions of whole blood cells were fixed with 2% PFA for 10 min. The cell suspension was dropped onto glass slides and air dried overnight. Then the single cells were ready for the following staining procedure. The single cells were permeabilized (0.1% Triton X-100, 0.1M Glycine, PBS) for 10 min. slides were washed and blocked with blocking buffer (3%BSA + 0.1% Tween-20 + 10% normal donkey serum) for 1 hour. Incubations with primary antibodies were performed overnight at 4°C. Incubation with secondary antibodies conjugated to Alexa Fluor (Jackson Immuno Research) was performed for 1.5 hours. After several washes with PBS, sections were stained with DAPI for 5 min at RT, rinsed with water and mounted with PermaFluor mounting medium (Beyotime). For human samples, The isolated neutrophils were fixed in 2% paraformaldehyde (PFA) and smears were prepared for immunofluorescence analysis. The slides were washed and blocked with blocking buffer (3%BSA + 0.1% Tween-20 + 10% normal donkey serum) for 1 hour. Slides were then stained with primary antibodies (mouse anti-CD15, Abcam, ab241552; rabbit anti-STEAP4, Proteintech,11944-1-AP) overnight at 4°C. Next, secondary antibodies conjugated to Alexa Fluor (Jackson Immuno Research) was performed for 1.5 hours. After several washes with PBST, sections were stained with DAPI for 5 min at RT, rinsed with water and mounted with PermaFluor mounting medium (Beyotime). Fluorescence images were acquired using a confocal microscope (Olympus FV4000). All the images were further processed with OlyVia software.

### Single-cell RNA-seq data processing and unsupervised clustering

scRNA-seq data were aligned and quantified using the CellRanger toolkit v7.1.0 against the reference genome mm10. Preliminary filtered data generated from Cell Ranger were then imported in the R-based package, Seurat v4.3.0 for downstream analysis (Hao et al., 2021). The high-quality cells were kept if satisfied the two metrics: (1) >= 200 expressed genes; >= 800 UMIs and mitochondrial UMIs <= 30% (2) <= 6000 expressed genes and <= 40000 UMIs. Scrublet was then applied to remove potential doublets with an expected doublet rate of 6%, and cells with more than 0.95 quantiles of doublet score were removed in the downstream analysis (Wolock et al., 2019). The raw count data are normalized using scale factor 10^4^ by the NormalizeData function. The top 2000 highly variable genes were selected by *FindVariableFeatures* (selection.method=”vst”). The scaled gene expression levels of these genes were used for principal component analysis (PCA) for dimensionality reduction using *RunPCA* function with parameter “npcs=30”. To remove batch effects between samples, we performed the Harmony algorithm to generate batch-corrected principal components (PC) using *RunHarmony* function with group.by.vars=”sample”. Next, the top 20 PCs were used for graph-based clustering by *FindNeighbors* and *FindClusters* functions. All cells were projected to a two-dimension plot for visualization based on the Uniform Manifold Approximation and Projection (UMAP) method implemented by the *RunUMAP* function using the top 20 PCs.

### Cell cluster annotation

The major cell types were annotated based on the first-round clustering according to well-known marker genes: *Epcam, Krt19, Cdh1* for epithelial cells, *Pecam, Cldn5* for endothelial cells, *Col1a1*, and *Msln* for stromal cells. The immune cells are marked by *Ptprc*, and then clustered based on *Cd3d*, *Cd3e, Cd4, Cd8a for T cells, Nkg7, Ncr1* for natural killer cells, *Cd79a* for B cells, *Ly6c2, Fcer1g, Cd68, Marco, Csf1r, Csf3r, Cd209a, Lamp3, Xcr1*, *Fscn1, S100a8* for myeloid lineages. Next, the second-round unsupervised clustering was based on each major cell type (T and NK, myeloid cells, epithelial, endothelial, stromal cells) by the same procedure with the first-round clustering. Different resolution parameters for unsupervised clustering were examined. The markable genes of each cell cluster are identified by the *FindAllMarkers* function in Seurat. Cell types were manually annotated according to the markers. Neutrophil marker genes in Fig. 2 are listed in Table S6.

### Composition analysis

To compare changes in neutrophil heterogeneity across normal lungs, primary lung tumors and metastatic lungs (Fig. 2), we first calculated proportion of each neutrophil subset within total neutrophils in each sample. Log2 fold changes were calculated by comparing the mean cell proportion in metastatic tumor samples against corresponding means in either normal controls or primary tumors. Statistical significance was determined using Wilcoxon-Mann-Whitney tests (two-group comparisons: met. Vs normal, met. Vs pri.) implemented through the wilcox_test function in R package coin, with the false discovery rates control via Benjamini-Hochberg adjustment. Metastasis-enriched neutrophil subsets with FDR<0.05 and log2FC>1 were highlighted in the figure.

### Identification of maturity of neutrophils

To predict the cell differentiation order among neutrophil cell states, neutrotime signature genes published by Grieshaber-Bouyer and collegues were applied using calculate signature score (Grieshaber-Bouyer et al., 2021). N1 cluster was highly expressed immature neutrophils markers (Xie et al., 2020), such as *Camp, Ngp, and Ltf*, and exhibited earliest Neutrotime. Therefore, N1 cluster was identified as immature neutrophils and chosen as the initiated state in the following trajectory analysis.

### Trajectory analysis of neutrophil subsets

To predict the cell differentiation trajectory, we applied RNA velocity and Slingshot in Fig. 2G. RNA velocity is performed by the following steps: First, the spliced and unspliced UMIs were distinguished by the python-based velocyto tool. Next, the data was imported into the scVelo package to estimate RNA velocities. After filtering the genes and normalized data by the scvelo.pp.filter_and_normalize function with default parameters, the moments were calculated by scvelo.pp.moments. The generalized dynamical model was used to estimate RNA velocity using function scvelo.tl.recover_dynamics, scv.tl.velocity(mode=”dynamical”) and scv.tl.velocity_graph. Finally, the velocities were projected onto the UMAP embedding using scv.pl.velocity_embedding_stream function for visualization (Fig 2G). Slingshot R package(version 2.0.0) was used to identify the neutrophil lineage structures. Slingshot was implemented in the analysis after dimensionality reduction and clustering of the integrated neutrophil data. UMAP embedding and cell cluster labels served as an input to the function slingshot (reducedDim, clusterlabels, start.clus = “N1”, extend= “n”, stretch = 0, omega = FALSE). The global lineage structure was identified with a cluster-based minimum spanning tree (MST), and then principal curves were fitted to model the development of cells along each lineage.

### Identification of the MNsig signature genes

The MNsig neutrophil signature genes were obtained from the differentially expressed genes in the MN_L_ subset versus other subsets using the following threshold: 1) log_2_ fold change > 0.4 and adjusted p-value <0.05 in the MN_L_. We obtained 65 genes used as MNsig (Table S4). Signature score for each cell of the integrated data across all conditions was calculated by *AddModuleScore* function in Seurat with default parameters, which estimates module activity based on average gene expression (Fig. 2). The identification of enriched Gene Ontology terms of MNsig genes were employed by *clusterProfiler* (Yu et al., 2012) and visualized by R package *GOplot* using chord diagram (Walter et al., 2015).

### Identify the highly-expressed membrane protein in the MN neutrophil subset

The Gene Ontology “plasma membrane” GO:0005886 were used to select membrane protein-coding genes. The MN_L_ up-regulated genes comparing to other neutrophil subsets (adjusted p-value < 0.05, expression percentage > 10% and log_2_ fold change >0.5, calculated from *FindAllMarkers* function described above) were used to select membrane proteins. Among these proteins *Steap4*, *Kit*, *Mif*, *Plscr1*, *Ltb4r1*, *Mcam* and *Vim*, *Steap4* has the highest expressing percentage (55%), and exhibiting specificity to lung metastasis.

### Screening for candidate membrane protein specific to lung metastasis

For each sample, we calculated the expression percentage of genes in the MNsig. The samples were stratified into three groups: neutrophils from normal lung tissues, from primary lung cancer in the KP model, and from lung metastases in the MMTV-PyMT model. For each gene, we calculated the mean expression percentage within each group, and determined the log2 fold changes (log2FC) when comparing MMTV-PyMT with either the normal, or KP groups. To address extreme log2FC valued resulting from small expression percentages, we introduced a “proportion-weighted log2FC” metric. This metric combines the mean expression percentage in the lung metastasis group with log2FC, providing a balanced measure that accounts for both the absolute value of expression percentage and the relative changes across comparisons. The proportion-weighted log2FC ensures the selection of genes with both high expression percentages and strong specificity for lung metastasis.

### Validation in public lung metastasis datasets

To validate the MNsig signature across diverse mouse models, we curated and re-analyzed several publicly available scRNA-seq and bulk RNA-seq datasets. These data includes: (1) the orthotopic 4T1 model and AT3 lung metastasis models from Gong et al., (GSE217143 and GSE217595) (Gong et al., 2023),. In this study, lung neutrophils were isolated via fluorescence-activated cell sorting (FACS) using the markers CD45^+^CD11b^+^Ly6C^low^Ly6G^+^ for subsequent scRNA-seq. (2) the orthotopic M3-9-M lung metastasis model from Kaczanowska et al., (Kaczanowska et al., 2021) (GSE168297), which profiled lungs at a pre-metastatic stage (day 15 post-inoculation). We re-processed this dataset and identified the neutrophil cluster based on high expression of canonical markers (Csf3r, Ly6g, S100a9) and the absence of lineage markers for macrophages (Csf1r), T-cells (Cd3d), and NK cells (Nkg7). (3) We also incorporated data from a genetically engineered mouse model of mammary tumor, the *K14*-Cre;*Cdh1*^fl/fl^;*Trp53^f^*^l/fl^ (KEP) mice from (Garner et al., 2025). This dataset included neutrophils isolated from the lung of wild-type and KEP mice using Ly6G^+^CD11b^+^ sorting and performed bulk RNA-seq sequencing. We performed differential gene expression analysis between the two groups, which confirmed the significant upregulation of both the MNsig signature and *Steap4* in lung metastasis (Fig. S2I and S3B).

### Prediction of lung metastasis using membrane protein genes on neutrophils based on scRNA-seq data

To evaluate the performance of STEAP4 and other known membrane proteins in predicting lung metastasis, we utilized the expression percentage of these proteins on neutorphils as indicators, based on scRNA-seq data. Using expression percentages, not gene expression values, could help mitigate biases that arise from batch effects across different datasets. The validation dataset included lung metastasis samples from our B16-F10 experimental metastasis model, as well as samples from 4T1 and AT3 models Gong and colleagues (Gong et al., 2023), and from M3-9-M model from Kaczanowska and colleagues (Kaczanowska et al., 2021). The control samples were integrated from neutorphils in the normal lungs from our data and the above published data. For each sample, the expression percentage of each gene in neutrophil was calculated by determining the frequency of non-zero expressing cells among neutrophils identified through scRNA-seq profiles. ROC analysis was conducted using the R package pROC, with the roc function for plotting ROC curves and the auc function for calculating the Area Under the Curve (AUC).

### Statistical analysis

Unless otherwise mentioned, the data are depicted as mean ± s.e.m. (as denoted in the figure legends). The figures showcase data gathered from numerous independent experiments, which were conducted on separate occasions using distinct samples. By default, the data displayed within the figure panels are derived from a minimum of three separate experiments. To assess the statistical significance of the mean differences between samples, two-tailed Student’s t-tests were employed.\**p*<0.05, \*\**p* < 0.01, \*\*\**p* < 0.001.

## Data and code availability

Sequencing data will be available through GEO and accession number will be provided by publication date. The data collected from public studies were listed in Supplementary Table S7. The code is available on our github website: https://github.com/AstreChen/lungmet.

## Authors’ contributions

C.A., H.W., L.G.N., and Z.Z. conceptualized this study. H.W. and C.A. designed the experiments. C.A., H.W., H.Y. wrote the original manuscript. L.G.N. and Z.Z. modified and reviewed the manuscript. H.W., H.Y., J.W, C.L. and L.Q.W performed experiments. C.A. performed computational analysis. Z.-H.Z., F.Z. and C.C organized and performed the experiments involving human samples. Q.G. and L.Z. assisted with the experiments and data analyses. M.S.F.N. contributed to the analysis of murine samples and revised the manuscript. C.A., H.W., L.G.N. and Z.Z. discussed and interpreted the data. Z.Z., L.G.N. and H.W. acquired fundings and administrated this project. Z.Z. supervised this project. All authors contributed to manuscript editing.

## Acknowledgments

This project was supported by funding from the National Key Research and Development Program of China (2023YFF1204700) (Z.Z.), National Natural Science Foundation of China (92374205) (L.G.N.), National Natural Science Foundation of China (82522001, 82470057) (H.W.), Shanghai Sailing Program (24YF2722400) (H.W.) and Shanghai Oriental Talent Program (QNWS2024087) (H.W.). Part of the analysis in this study was performed on the High Performance Computing Platform of the Center for Life Science (Peking University). We thank the Laboratory Animal Center of Peking University for advice and technical support. We thank Core facility of Basic Medical Science, Shanghai Jiao Tong University School of Medicine for technical support of flow cytometry sorting and imaging experiment. We thank members of the Zhang lab for their discussion and feedbacks on this study. We thank members of the Wu lab for experimental technical discussion and support. We thank all patients for participating in this study.

## Supplementary Tables S1-7

Supplementary Table S1. Clinical characteristics of human subjects analyzed by immunofluorescence staining.

Supplementary Table S2. Murine samples for scRNA-seq in this study.

Supplementary Table S3. T cell activation signatures, related to Fig. S1.

Supplementary Table S4. Metastasis-specific signature in neutrophils.

Supplementary Table S5. Clinical characteristics of human subjects providing fresh blood samples for flow cytometry analysis.

Supplementary Table S6. Marker genes for neutrophil subsets in scRNA-seq analysis.

Supplementary Table S7. The published data used in this paper.

## Supplemental material

**Figure S1.**
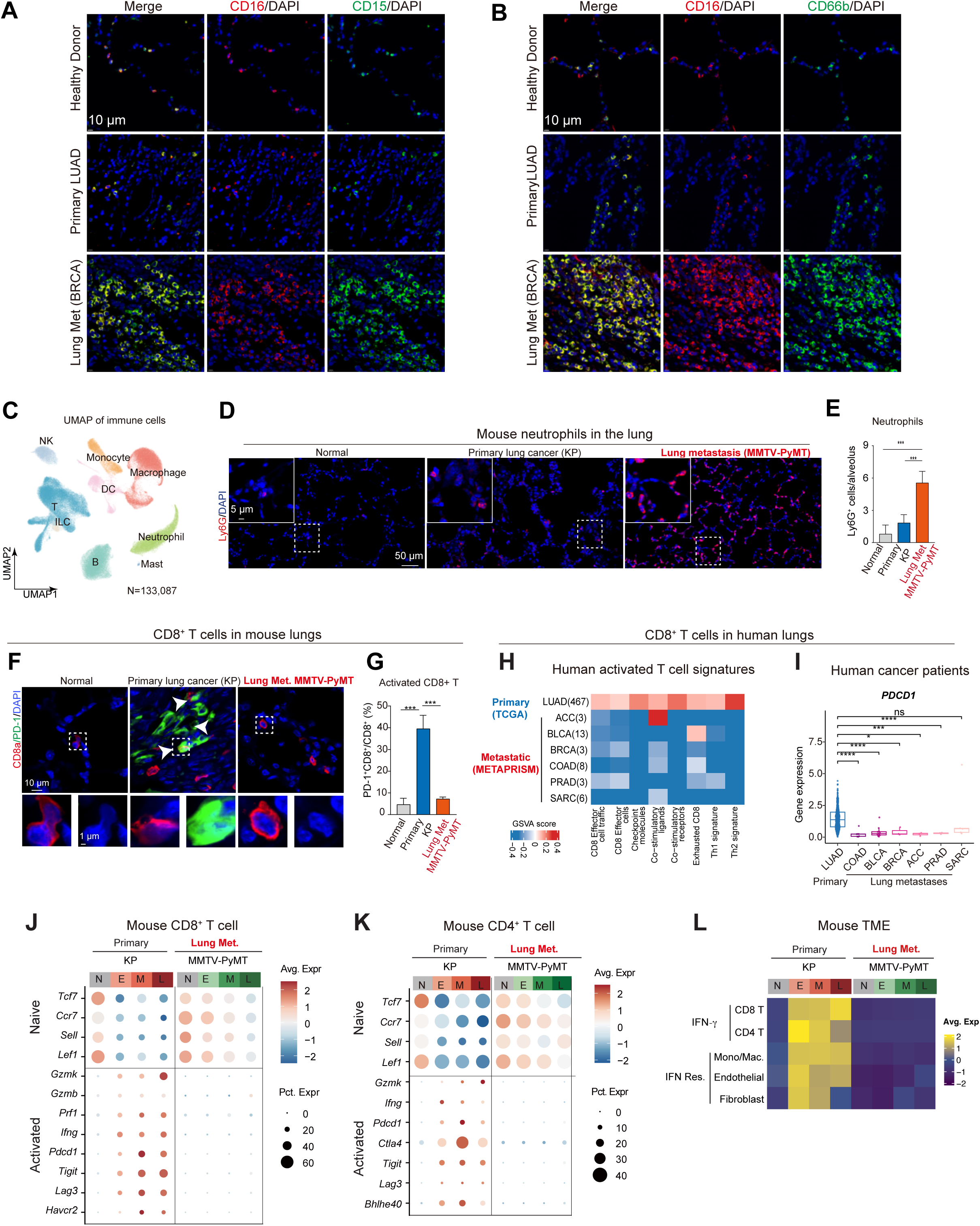
Immune phenotypes of lung metastases in human patients and mouse models. **A-B,** Immunofluorescence staining of neutrophils in the human lung tissues using CD15 and CD16 (A) or CD66b and CD16 (B). Scale bars, 10 μm. **C,** UMAP plot of immune cells from lung tissues of mice shown in MMTV-PyMT (15 mice) and KP models (13 mice). NK, natural killer cells. **D-E,** Neutrophils were stained with antibodies against Ly6G in the lungs from normal, KP, and MMTV-PyMT mice (D), and their number in alveolus was quantified (E). (Mean ± SEM, n = 3). Student’s t-test. \*\*\**p*<0.001. Scale bars, 50 μm and 5 μm. **F-G**, Tissues were stained with antibodies against CD8a and PD1, with white arrowheads indicating CD8^+^ PD1^+^ T cells (F). The percentages of CD8a^+^ PD1^+^ T cells were quantified in (G). (Mean ± SEM, n = 3). Student’s t-test. \*\*\**p*<0.001. Scale bars, 10 μm and 1 μm. **H,** Heatmap showing comparison of the expression scores of T cell activation signatures between lung primary tumors (LUAD, n=436) and metastatic tumors (n=100) based on RNA-seq data of TCGA and METAPRISM cohort. **I,** Box plots showing the expression level of *PDCD1* in the primary lung cancer patients and patients with lung metastasis. \**p*<0.05, \*\**p*<0.01, \*\*\**p*<0.001, \*\*\*\**p*<0.0001, ns, not significant, *p* values from Student’s t-test. **J-K,** Bubble plots showing activity of the functional genes in CD8^+^ T cells (J) CD4^+^ T cells (K) from primary KP lung tumor and metastatic lung tumors from MMTV-PyMT. *N* = normal, *E* = early, *M =* mid, *L* = late stage. **L,** Heatmap showing the scaled gene expression of *Ifng* in CD8^+^ and CD4^+^ T cells; and the signature score of response to interferons of GO terms in lung tissue samples from different time points of mouse models. *N* = normal, *E* = early, *M =* mid, *L* = late stage.

**Figure S2.**
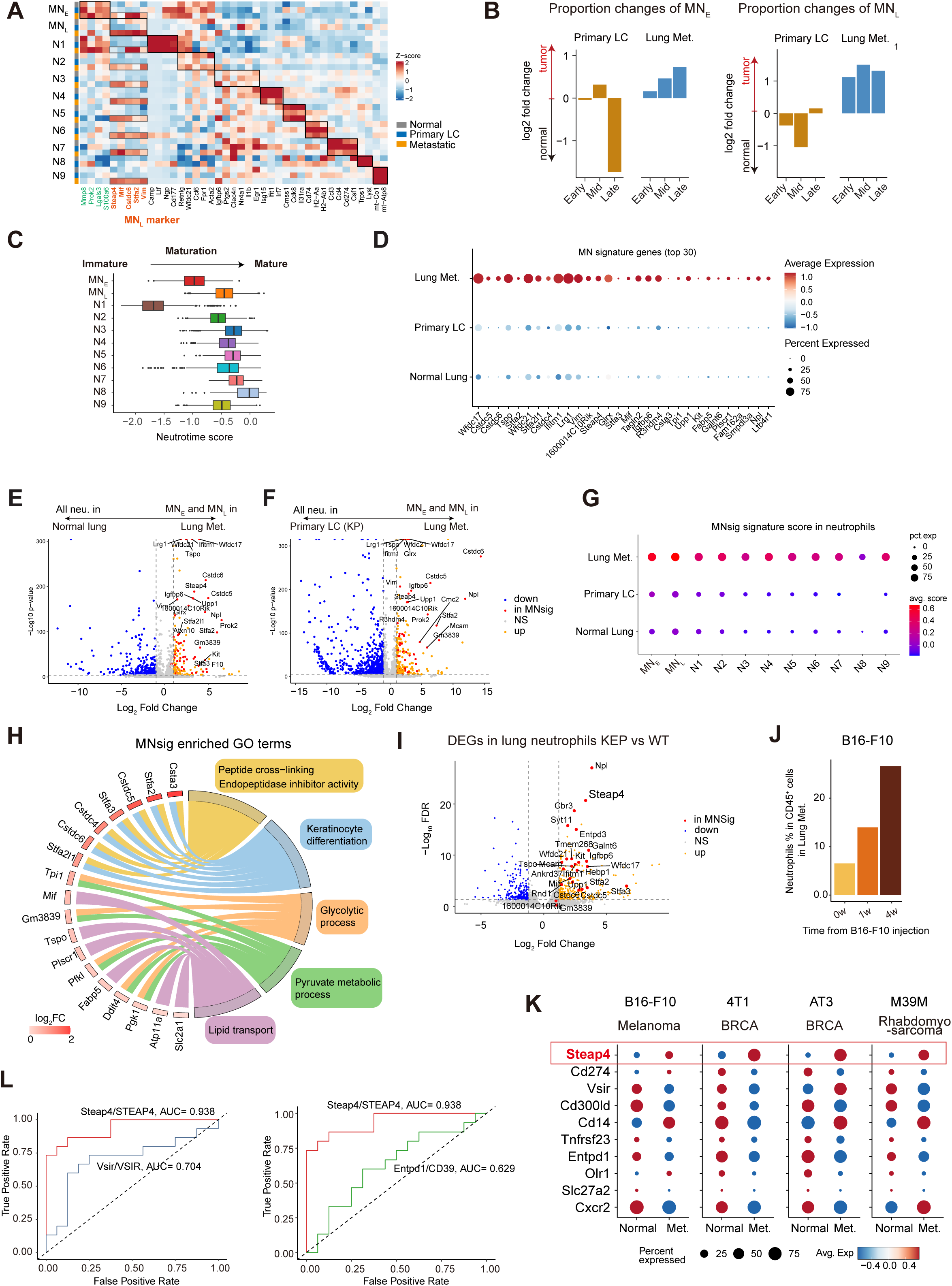
Characterization of neutrophils via scRNA-seq analysis. **A,** Heatmap showing top marker genes of each neutrophil subset split by conditions shown in Fig. 2A. See Table S6 for all top 50 marker genes. **B,** Bar plots showing changes in cell proportion of MN_E_ and MN_L_ subsets in primary lung tumors versus metastatic lung tumors via log fold change calculated based on the mean proportion across samples in each group. **C,** Neutrotime signature score of each subset of neutrophil in the normal, KP and MMTV-PyMT lungs. The large neutrotime scores represent a mature state. **D,** Bubble plot depicting expression level and percentage of the MN signature genes in neutrophils from normal lungs, primary lung tumors(KP) and lung metastases (MMTV-PyMT). **E,** Volcano plot showing the differentially expressed genes between metastasis-specific neutrophils (MN_E_+MN_L_) in the MMTV-PyMT and neutrophils from normal lungs. **F,** Volcano plot showing the differentially expressed genes between metastasis-specific neutrophils (MN_E_+MN_L_) in the MMTV-PyMT and neutrophils in the primary lung tumor (KP model). **G,** Bubble plot depicting the level and percentage of MNsig score within different lung metastasis mouse models. Scores are calculated using AddModuleScore. **H,** Chord diagram showing MNsig genes and the enriched pathways in GO terms. The log2 Fold Change represents the expression change in MN_L_ versus other neutrophil subsets. **I,** Volcano plot displaying differential gene expression in lung neutrophils from KEP versus wild-type mice. The data were re-analyzed using the bulk RNA-seq dat from the study (Garner et al., 2025). **J,** Bar plot showing the percentage of neutrophils in the CD45^+^ cells harvest from lungs in the B16-F10 experimental metastasis model. **K,** Bubble plot depicting the level and percentage of the cell surface protein genes within different lung metastasis mouse models. **L,** The ROC curves showing the performance of the expression percentage of VISTA, CD39 and STEAP4.

**Figure S3.**
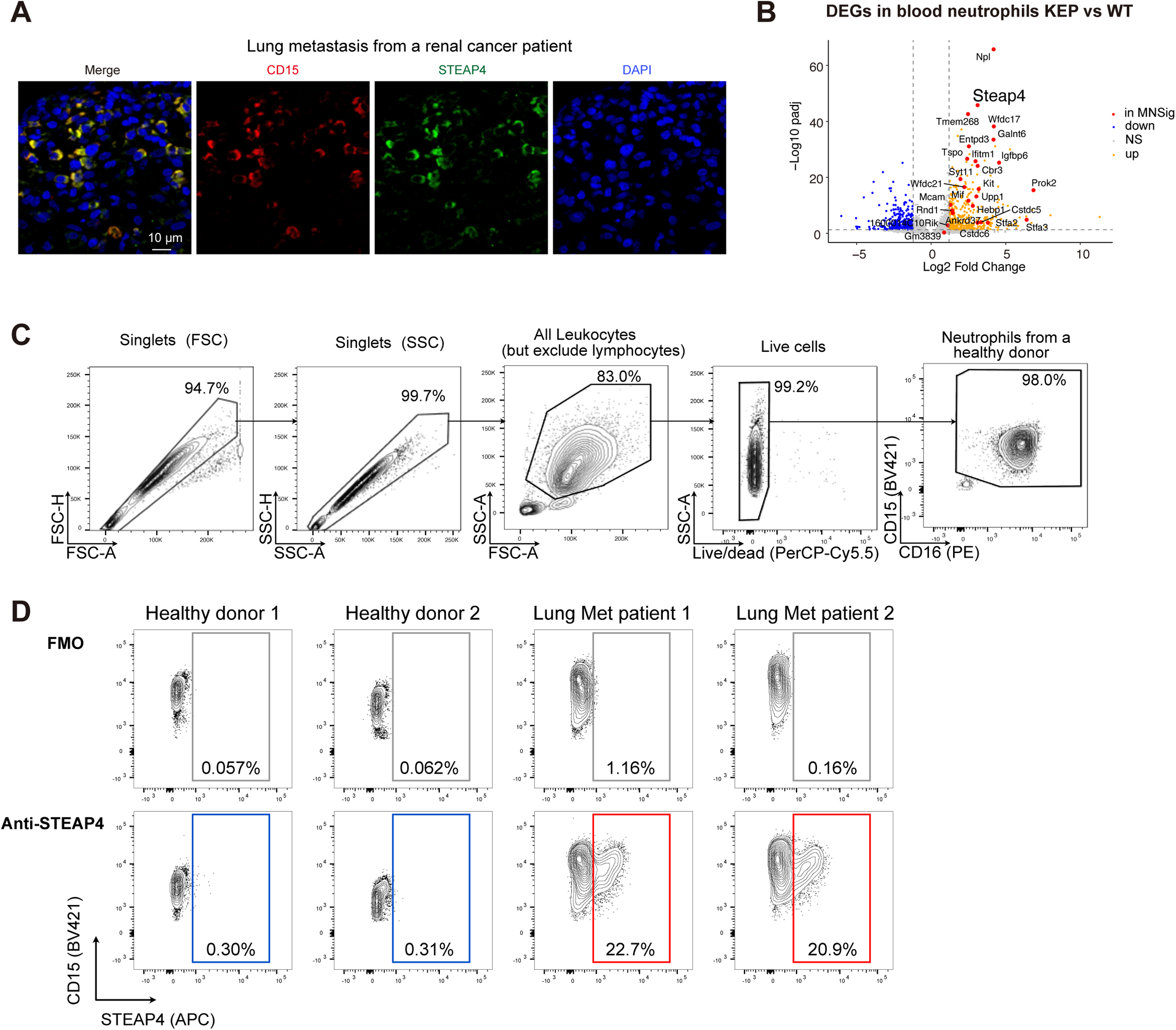
Confirmation of STEAP4^+^ neutrophils in the lung tissue and peripheral blood in human patients. **A,** Representative immunostaining of lungs from a renal cancer patient with lung metastases using antibodies agaist CD15 and STEAP4. Scale bars, 10 μm. **B,** Volcano plot displaying differential gene expression in blood neutrophils from KEP versus wild-type mice. The data were re-analyzed using the bulk RNA-seq data from (Garner et al., 2025). **C,** Gating strategies for selecting neutrophils from fresh blood are shown in Fig. 4E. **D,** Representative contour plots showing the frequencies of STEAP4^+^ neutrophils in the healthy donors and patients with metastasis, gated using the fluorescence minus one (FMO) control.

