## Supplemental Table S1 for "STEAP4^+^ neutrophils: a promising circulating biomarker for lung metastasis"

Supplementary Table S1. Clinical characteristics of human subjects analyzed by immunofluorescence staining.

| Group | Sex | Neutrophil in blood（%） | Primary cancer type | Application |
| --- | --- | --- | --- | --- |
| Donor | F | 47.1 | - | Immunofluorescence staining |
| Donor | M | 76 | - | Immunofluorescence staining |
| Donor | F | 47.1 | - | Immunofluorescence staining |
| Donor | M | 78.1 | - | Immunofluorescence staining |
| Donor | M | 63.2 | - | Immunofluorescence staining |
| Donor | F | 78.1 | - | Immunofluorescence staining |
| Donor | F | 68.6 | - | Immunofluorescence staining |
| Donor | F | 53.9 | - | Immunofluorescence staining |
| Donor | M | 67.3 | - | Immunofluorescence staining |
| Donor | M | 45.2 | - | Immunofluorescence staining |
| LUAD | F | 62 | Lung adenocarcinoma | Immunofluorescence staining |
| LUAD | F | 59.9 | Lung adenocarcinoma | Immunofluorescence staining |
| LUAD | F | 51.5 | Lung adenocarcinoma | Immunofluorescence staining |
| LUAD | F | 64.6 | Lung adenocarcinoma | Immunofluorescence staining |
| LUAD | M | 61.2 | Lung adenocarcinoma | Immunofluorescence staining |
| LUAD | M | 60.3 | Lung adenocarcinoma | Immunofluorescence staining |
| LUAD | M | 53.1 | Lung adenocarcinoma | Immunofluorescence staining |
| LUAD | F | 65.9 | Lung adenocarcinoma | Immunofluorescence staining |
| LUAD | F | 72.3 | Lung adenocarcinoma | Immunofluorescence staining |
| LUAD | F | 60.3 | Lung adenocarcinoma | Immunofluorescence staining |
| Lung metastasis | F | 56.9 | Breast cancer | Immunofluorescence staining |
| Lung metastasis | M | 71.4 | Kidney cancer | Immunofluorescence staining |
| Lung metastasis | F | 49.2 | Breast cancer | Immunofluorescence staining |
| Lung metastasis | M | 61.7 | Liver cancer | Immunofluorescence staining |
| Lung metastasis | F | 53.1 | Breast cancer | Immunofluorescence staining |
| Lung metastasis | F | 34.6 | Intestinal cancer | Immunofluorescence staining |
| Lung metastasis | M | 59.3 | Intestinal cancer | Immunofluorescence staining |
| Lung metastasis | M | 67.7 | Colon cancer | Immunofluorescence staining |
| Lung metastasis | M | 68.5 | Intestinal cancer | Immunofluorescence staining |
